# Topography structures of arthropod communities revealed by leaf-derived environmental DNA on O’ahu, Hawai’i

**DOI:** 10.64898/2026.04.09.716978

**Authors:** Sven Weber, Leke Hutchins, Pritam Banerjee, Willow Callaghan, Alex Augustus Farrow, Jeremy Andersen, Rosemary Gillespie, George Roderick

## Abstract

Arthropod communities on oceanic islands are shaped by spatial isolation, environmental gradients, and biological invasions, yet their structure remains difficult to resolve due to incomplete taxonomic coverage. In particular, it remains unclear (i) how non-native arthropods can influence community composition and (ii) how they interact with native and non-native plants. To answer the first question, we used leaf-derived environmental DNA (eDNA) to characterize arthropod communities across elevational gradients on five ridges on O’ahu, using the native tree *Metrosideros polymorpha* as a standardized plant. To understand the second question, we compared leaf-derived eDNA from *Metrosideros polymorpha* (Native), *Acacia koa* (native), and *Psidium cattleianum* (invasive), co-occurring in two ridges on O’ahu. Additionally, to overcome limitations of reference databases, we applied NIClassify to infer native versus introduced status without requiring species-level identification. Across 96 leaf samples (with 851 Arthropod ASVs), we found arthropod richness increased with elevation, while the proportion of introduced taxa declined significantly. Community composition was primarily structured by ridge, with strong distance–decay relationships indicating high spatial turnover in both native and non-native assemblages. In contrast, plant species effects were context dependent and did not show a consistent native versus invasive signal. Threshold analyses identified a community transition (native vs introduced) near 500 m elevation. These results show that plant-derived eDNA can resolve spatial and environmental structuring of arthropod communities while capturing invasion dynamics under incomplete taxonomic knowledge. Classifier-based inference enables community-level ecological interpretation beyond reference-limited taxa, providing a scalable framework for biodiversity monitoring in data-poor systems.

## 1. Introduction

Arthropod communities are structured by the combined effects of mechanisms, including environmental filtering, dispersal limitation, species interactions, and evolutionary history (Chase & Leibold, 2009; Vellend, 2010). On oceanic islands, these processes can generate highly distinctive assemblages marked by endemism, ecological specialization, and strong turnover across heterogeneous habitats (Losos & Ricklefs, 2009; Whittaker & Fernández-Palacios, 2007). In Hawai i, one of the most remote archipelagoes on earth, arthropod communities have been shaped by both adaptive diversification and vulnerability to environmental change, producing faunas that are ecologically distinct but still incompletely documented (Graham et al., 2017). Because arthropods play central roles as herbivores, predators, decomposers, and pollinators, understanding how their communities are shaped over space and time is essential for evaluating ecological change in Hawaiian forests and in predicting response to environmental stressors.

Elevation gradients provide a powerful framework for studying mechanisms driving community structure because they integrate shifts in temperature, rainfall, vegetation, disturbance, and invasion pressure across relatively short distances. In Hawai i, lower elevations are associated with more disturbance and greater pressure from invaders, whereas higher elevations retain cooler conditions and a greater proportion of native vegetation (Loope et al., 2013; Vitousek et al., 1995). Such environmental gradients are therefore expected to influence not only overall community composition, but also the balance between native and introduced lineages. If invasion pressure is strongest at low elevations, arthropod communities are expected to be dominated by non-native species and be more homogeneous in these habitats, and become increasingly differentiated toward higher elevations, where native forest conditions persist and introduced taxa decline.

A major challenge in testing the of mechanisms that structure arthropod communities across elevational gradients is plant diversity, which can strongly influence associated arthropod assemblages. Plant species differ in architecture, chemistry, and phenology, and these differences can affect which arthropods colonize, persist, or feed on them (Novotny et al., 2002; Schaffers et al., 2008; Southwood, 1961; Strong et al., 1984; Tobisch et al., 2023). In Hawaiian forests, O’hi’a lehua, *Metrosideros polymorpha*, is especially well suited for standardized comparisons across habitat types because it is a foundational native tree that spans broad elevational and ecological ranges and forms the structural backbone of many wet and mesic forests (Barton et al., 2021; Percy et al., 2008). Focusing on O’hi’a therefore, allows arthropod communities to be compared across elevation while minimizing confounding variation caused by changing plant identity.

At the same time, plant identity remains biologically important and may explain additional local turnover of arthropod species beyond elevational effects alone (Di Marco et al., 2023). Native and introduced trees differ in the arthropod communities they support, both because of their traits and because of their ecological history within Hawaiian forests (Sax & Gaines, 2008). On O’ahu, the native koa, *Acacia koa*, and the invasive strawberry guava, *Psidium cattleianum*, often co-occur with O’hi’a on the same slopes, creating a useful contrast among native and introduced plants under comparable environmental conditions. These comparisons allow one to ask whether plant identity adds explanatory power beyond elevation, and the extent to which the relative representation of introduced versus native arthropods is associated simply with plant-origin pattern or instead with geographic factors (Denslow et al., 2024; Ellshoff et al., 1995; Gruner, 2004).

Many Hawaiian arthropods remain undescribed or lack robust DNA barcode representation, which limits species-level identification and complicates ecological interpretation (Porter & Hajibabaei, 2018b). Environmental DNA (eDNA) metabarcoding can recover diverse arthropod communities associated with plants, making large-scale comparisons possible, but the ecological associations of detected lineages often depend on whether they can be linked to known taxa and distributions (Banerjee et al., 2022, 2026; Krehenwinkel et al., 2022). In addition, arthropod DNA metabarcoding datasets often suffer from incomplete taxonomic and reference-sequence coverage, which obstructs ecological interpretation (Porter & Hajibabaei, 2018a; Weigand et al., 2019). In invasion ecology, this creates a specific conundrum: biogeographic status is often most interesting precisely for the taxa that are hardest to identify confidently to species.

To address this gap, NIClassify (https://github.com/tokebe/niclassify) was developed to allow inference of native versus introduced status from DNA sequence-derived features without requiring species-level BLAST identification. Rather than depending entirely on named matches in incomplete databases, this approach uses phylogenetic and distribution-linked signals to classify lineages as native (N) or introduced (I), even when taxonomic resolution is limited (Andersen et al., 2019). In principle, this allows invasion-status analyses to extend beyond the subset of taxa that can be assigned confidently by conventional DNA reference-based workflows, making it especially useful for diverse Hawaiian arthropod communities, for which incomplete reference coverage remains a major constraint.

Here, we set out to determine the role of geography and plant species in shaping the shifting balance of arthropod composition, using leaf-derived eDNA to characterize arthropod communities across elevational gradients in Hawaiian forests on multiple ridges, with O’hi’a as a standardized plant and additional comparisons among co-occurring koa and strawberry guava. We first evaluated the use of NIClassify to infer native versus introduced status from DNA sequence-derived features without requiring species-level identification through NCBI, BLAST, making invasion-status inference possible under incomplete Hawaiian reference coverage. Next, we tested predictions from two hypotheses to explain the relative abundance of introduced and native arthropod species across replicate elevational gradients: **(1)** When arthropod communities are standardized by sampling on O’hi’a, community differentiation (native vs introduced) will increase toward higher elevations, consistent with stronger native arthropod-native plant associations in less invaded, upper-elevation forests compared to more homogenization at lower elevations. **(2)** Plant identity, and especially plant origin (native or introduced), contributes additionally to local species turnover on shared slopes. Finally, we sought evidence for a change point in community structure with elevation, where arthropod communities change from dominated by introduced species to native species with elevation.

## 2. Material and Methods

### 2.1 Study design and Sampling

In Hawaii, *Metrosideros polymorpha* (O’hi’a) occurs across much of the Hawaiian archipelago but becomes particularly abundant on O’ahu from approximately 450 m elevation upward, where it often forms distinct forest stands (**Figure 1**). Sampling plots were therefore centered on mature *M. polymorpha* trees along replicated elevation gradients, selecting locations where leaf material could be collected from the ground without climbing. *Acacia koa* is primarily associated with mesic to wet montane forests at mid-elevations (Baker, 2009), while the invasive *Psidium cattleianum* is broadly distributed across low to mid elevations, where it readily colonizes disturbed habitats and can form dense stands from sea level to approximately 1,200 m, with its upper limit constrained by temperature and moisture gradients (Denslow et al., 2009, 2024).

**Figure 1.**
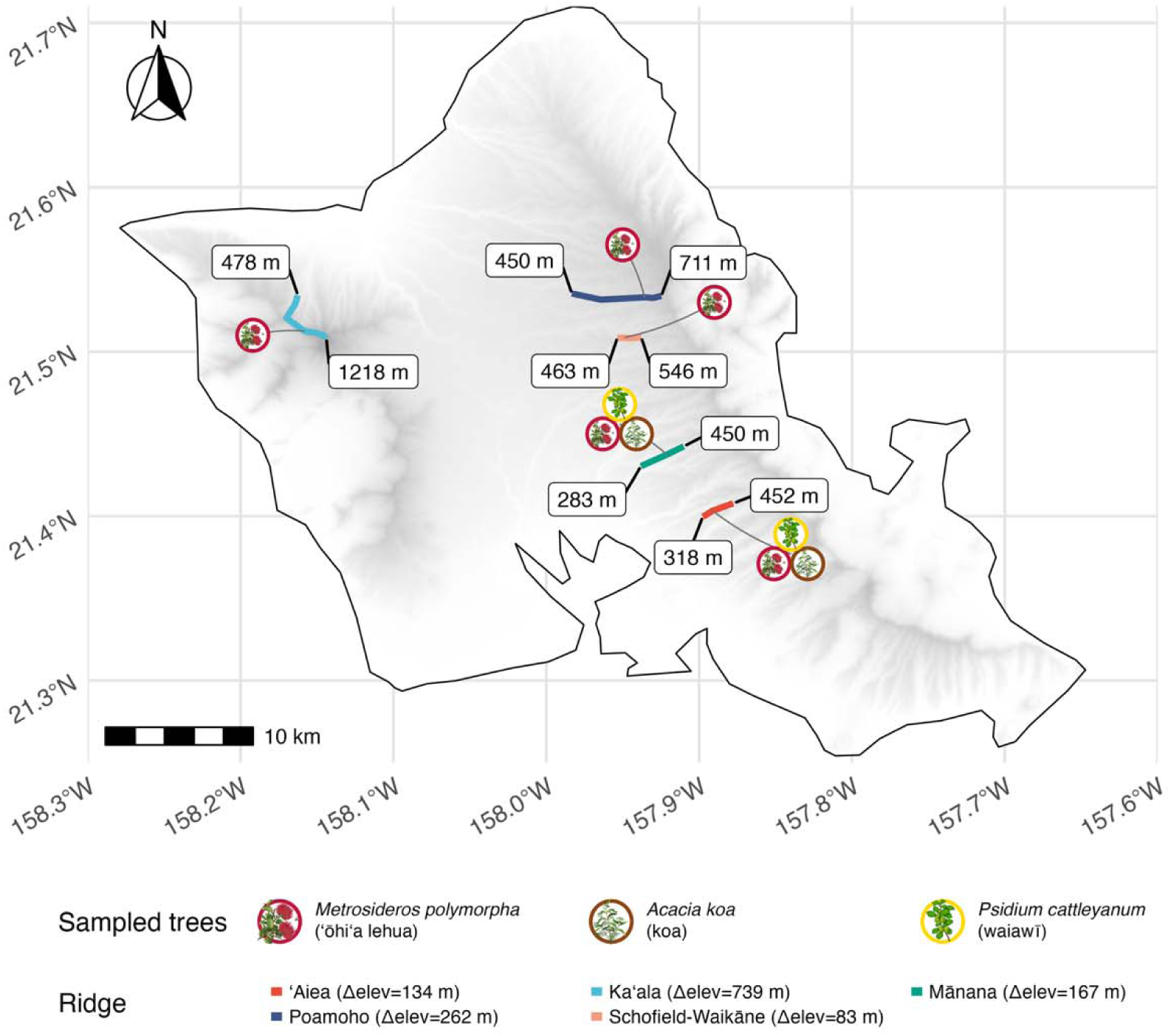
Map showing sampling approaches for leaf-derived eDNA from native and non-native plants on O’ahu, Hawai’i. Leaves from *M. polymorpha* (across all five ridges), *A. koa*, and *P. cattleianum* (from Aiea and Manana), were collected for plant-arthropod interactions analysis with eDNA. Labels indicate the minimum and maximum elevation of each ridge transect (in meters).

The range of elevations differed among ridges. Thus, we sampled the gradients for each location with the aim of getting samples for dividing roughly into low, mid, and high elevations. Leaf material was collected using single-use nitrile gloves (USA Scientific, USA) and disposable collection bags to minimize cross-contamination. We sampled ∼25 g of leaves per species (all three plants) from three separate plant individuals at each location (low, middle, high), from Aiea (318m – 452m) and Manana (283m – 450m) across both transects. A similar approach was applied for Ka ala (478m – 1218m), Poamoho (450m – 711m), and Schofield-Waikāne (463m – 546m), but collecting for *M. polymorpha*. Altogether, we collected leaves from only *M. polymorpha* across three different ridges (Ka ala, Poamoho, and Schofield-Waikāne), and *M. polymorpha*, *A. koa,* and *P. cattleianum* from two ridges ( Aiea and Manana) (**Figure 1**). Negative field controls, including open tubes exposed to ambient air and empty paper bags, were processed alongside all samples. Sampling details, with locations, elevations, and distribution, can be found in **Table S1**.We used our samples in two different ways to test our hypothesis: *M. polymorpha* from all five ridges was used to test the first hypothesis (H1), and *M. polymorpha* along with *A. koa* and *P. cattleianum* from two ridges was used to test the second hypothesis (H2) (**Figure 1**).

### 2.2. Leaf sample processing

Samples were returned to the field station and freeze-dried for at least 48 h using a Labconco FreeZone® 1 Liter Benchtop system (Model #7740020, Labconco, USA). Particularly wet samples required up to approximately 100 h to dry completely. After drying, we combined the three sub-samples (from three different individuals of same plants species, collected from the same locations) and homogenized using a household blender (Ninja®), following (Weber et al., 2024). Thus, 96 foliage eDNA samples collapsed to 32 biological samples. Homogenized material was transferred into one or two 50 mL Falcon tubes for long-term storage before DNA extraction (Stothut et al., 2024). To minimize carryover contamination, blender components were cleaned after each homogenization using a three-step decontamination protocol (Buchner et al., 2021).

### 2.3 DNA extraction, PCR, and Next Generation Sequencing

After homogenization, two replicates of 150 mg of powdered leaves were used for eDNA extraction using a CTAB-based protocol optimized for the reduction of PCR inhibitors (Weber et al., 2024). We used three PCR replicates (with an inline barcode combination) (Faircloth & Glenn, 2012) for each sample to maintain the reproducibility and recover the rare taxa. In total, we have two replicates of extraction and three replicates of PCR for each extraction, we have six technical replicates per sample. We used a two-step PCR approach, in first PCR an amplification of the genetic marker was amplified, followed by an indexing PCR to attach the Illumina sequencing adapters (Krehenwinkel et al., 2019). We chose a 180bp mitochondrial COI barcoding region developed by Jusino et al. (2019), where the forward primer was LCO1490 5′ GGTCAACAAATCATAAAGATATTGG 3′ (Folmer et al., 1994) and the reverse primer was CO1 CFMRa 5′ GGWACTAATCAATTTCCAAATCC 3′ (Jusino et al., 2019). Each 10 µL reaction contained 5 µL of 2× QIAGEN Multiplex PCR Master Mix (Qiagen, Hilden, Germany), 1 µL of DNA template, 0.5 µL of each 10 µM forward and reverse primer, 1 µL of Q-solution, and 2 µL of nuclease-free water. Thermal cycling conditions were 95 °C for 15 min, followed by 35 cycles of 94 °C for 30 s, 46 °C for 90 s, and 72 °C for 90 s, with a final extension at 72 °C for 10 min. PCR products were visualized on 1.5% agarose gels stained with GelRed (Fisher Scientific, USA). Then we pooled the triplicates together based on relative band intensity from the gel images. Afterwards, we performed a second PCR (indexing PCR) to attach Illumina sequencing adapters and 8-bp dual indexes to each sample, ensuring a minimum two-base difference between index sequences following the protocol of Lange et al. (2014). Reaction composition and cycling conditions were identical to the first PCR, except that only five amplification cycles were performed. Field blanks, filtration blanks, extraction blanks, and PCR blanks were included as negative controls throughout the workflow. Following final inspection by agarose gel electrophoresis, all PCR products were purified with 1.5× SPRI beads (AMPure XP, USA). Quality of all pools was assessed using a Qubit 3.0 fluorometer (Invitrogen, Thermo Fisher Scientific, USA) and a 2100 Bioanalyzer (Agilent Technologies, USA), then pooled in equal amounts into a single tube. All samples (including negative controls) were sequenced (targeting ∼50,000 reads per sample) on an Illumina NextSeq platform using P2 chemistry (2x300 bp), with 30% Phix, including all negative controls for each step sequenced alongside field blanks as explained above at QB3 Genomics Facility, University of California, Berkeley.

### 2.4 Bioinformatic analysis

The samples were demultiplexed using DRAGEN v3.10 (Illumina Inc., San Diego, California, USA) with no mismatches allowed. Demultiplexed fastq files were merged using PEAR (Zhang et al., 2014) with a minimum overlap of 50 and a minimum phred quality score of 20. The merged reads were additionally filtered for a minimum quality of Q33 over >90% of the sequence and then transformed to fasta files using the FASTX-Toolkit. PCR primer sequences were then trimmed off from the merged reads using a custom UNIX script, allowing degenerate bases to vary in the search patterns and acknowledge the provided inline barcodes for the sample replicates. The processed reads were dereplicated and clustered into ASVs using VSEARCH (Rognes et al., 2016). The minimum size cluster was determined with eight occurrences and combined with a de novo chimera removal.

All resulting ASVs were searched against the NCBI database using BLASTn (updated 02/2025) with a maximum of 10 target sequences (Altschul et al., 1990). Taxonomy was then assigned to the resulting BLAST output using a custom python script (Schöneberg, 2023). The ASV table was then constructed for all samples using VSEARCH (Rognes et al., 2016) (**Supplementary Material S3**). All PCR replicated were merged after checking their correlation with a similarity of over 75%. We filtered out all hits below 90% sequence similarity, and to maintain the focus of this study on plant-arthropod interactions, all non-arthropod ASVs were removed. All hits below 90% similarity to a reference were omitted. Order level was filtered to >93%, family level was filtered to >95%, and species level was filtered to >98% reference hit to the database to account for the small fragment size of the amplicon (Krehenwinkel et al., 2022).

### 2.5 Invasion status classification

Native and non-native status of all ASVs (>90%) was predicted using the program NIClassify (https://github.com/tokebe/niclassify). We used a curated Hawaiian arthropod checklist containing more than 12,000 species entries and 3,064 labeled ASVs as training data (**<u>Supplementary Material S4</u>**). ASVs were first assigned to order-level groups, and then NIClassify calculated order-specific phylogenetic features based on local clustering structure and sequence divergence to predict native/non-native status (Andersen et al., 2019). Models were trained with a random forest classifier using 1,000 trees after removing any overlap between training and prediction sets. For downstream analyses, ASVs were classified as introduced when the predicted probability of introduced status was at least 0.51 (**Supplementary Material 2**). Additionally, assigned taxa were checked manually against curated Hawaiian sources, and ambiguous matches were labeled uncertain. We compared NIClassify assignments with BLAST-only and IBIS-based lookups as sensitivity checks, but all main analyses in this study are based on NIClassify probabilities because of limitations in the Hawaiian reference database, and the alternative approaches either over-assigned introduced status or remained unresolved under sparse references.

### 2.6 Statistical analysis

All analyses were performed in R 4.4.3 (**R version 4.4.3 (Trophy Case)**) using the tidyverse (Wickham et al., 2019), patchwork data wrangling for visualization and ggpubr (Kassambara, 2018) for incorporating statistical values and test into the plots. Linear regressions of richness and Shannon diversity against elevation were run for context but are not emphasized (Oksanen et al., 2013). Additional site-level contrasts (ridge vs elevation band, native vs introduced fractions, extraction volume comparisons) were tested with linear mixed models (*lme4*) and non-parametric rank tests (Kruskal–Wallis with Dunn’s post hoc comparisons; *emmeans*). Overlap in taxa among tree species and elevation bands was visualized with Venn/UpSet diagrams (*ComplexUpset* (Krassowski, 2021). Jenks natural breaks were used to define elevation and rainfall bands (*classInt* (Bivand et al., 2015)

#### 2.6.1 Diversity, community composition, and Invasion pattern

We calculated Bray–Curtis dissimilarities from ASV tables and visualized community structure using non-metric multidimensional scaling (metamds() with k=2, trymax = 1337 from *vegan*). Effects of elevation, site, and tree species identity were tested with permutational multivariate analysis of variance (PERMANOVA; *adonis2* in *vegan)*, using 999 permutations and reduced models to isolate main effects. To evaluate group heterogeneity, we tested multivariate dispersion with *betadisper* (*vegan*). We further partitioned Sørensen dissimilarity into turnover and nestedness components (*betapart*) and assessed distance–decay relationships between community similarity and geographic distance using haversine distances (*geosphere*) with Mantel tests (*vegan*).

Species accumulation curves, raw and rarefied richness were used to compare diversity across elevation bands and tree species (*vegan*). To test whether the proportion of introduced ASVs varied along gradients, we applied binomial generalized estimating equations (GEEs; *geepack*) with ridge as the clustering variable and elevation band (low, mid, high) as predictor.

#### 2.6.2 Change-point analyses

To identify taxon-specific and community-level responses along elevation, we used Threshold Indicator Taxa Analysis (TITAN2; Baker et al., 2015). Analyses were performed on a presence/absence matrix (taxa occurring in ≥ 3 times), with elevation (m) as the gradient. We used minSplt = 5, numPerm = 250, boot = TRUE, nBoot = 500, imax = TRUE, pur.cut = 0.95, and rel.cut = 0.95. Community-level change-points were summarized using sum(z) (filtered/unfiltered), bootstrap medians, and 95% quantiles, and visualized as density and violin plots (ggplot2). Taxon-level breakpoints were extracted for taxa meeting purity/reliability criteria, with responses classified as z (increasing with elevation) or z (decreasing). We also evaluated sensitivity using presence/absence data and visualized sampling density across gradients.

## 3. Results

We analyzed 96 leaf eDNA samples collapsed to 32 samples representing five ridges on the island of Oahu, detecting 851 arthropod ASVs spanning 26 orders. The most abundant orders by read share were Lepidoptera (12.65%), Hemiptera (12.57%), and Acari (9.02%). In total, we kept 1,071,406 reads (mean ± SE: 33,481 ± 4,696 reads per sample) after filtering for arthropod reads. Mean ASV richness was 36.18 ± 3.98 per biological sample.

### 3.1 NIClassify enables invasion-status inference under incomplete reference coverage

Native and introduced status distributions differed significantly among BOLD, IBIS, NCBI, and NIClassify (χ² = 118.51, df = 3, p < 2.2 × 10 ¹; **Supplementary Material 1**; **Figures S1–S2**; **Table 1**). Most of the reference-based identification methods (BOLD, IBIS, NCBI) could not resolve the status of arthropods when they were identified above or similar to family level **(Figure 2**). As incomplete reference databases can influence the final results, we selected NIClassify to predict the arthropod native and introduced status. Thus, all downstream status analyses are therefore based on NIClassify assignments.

**Figure 2.**
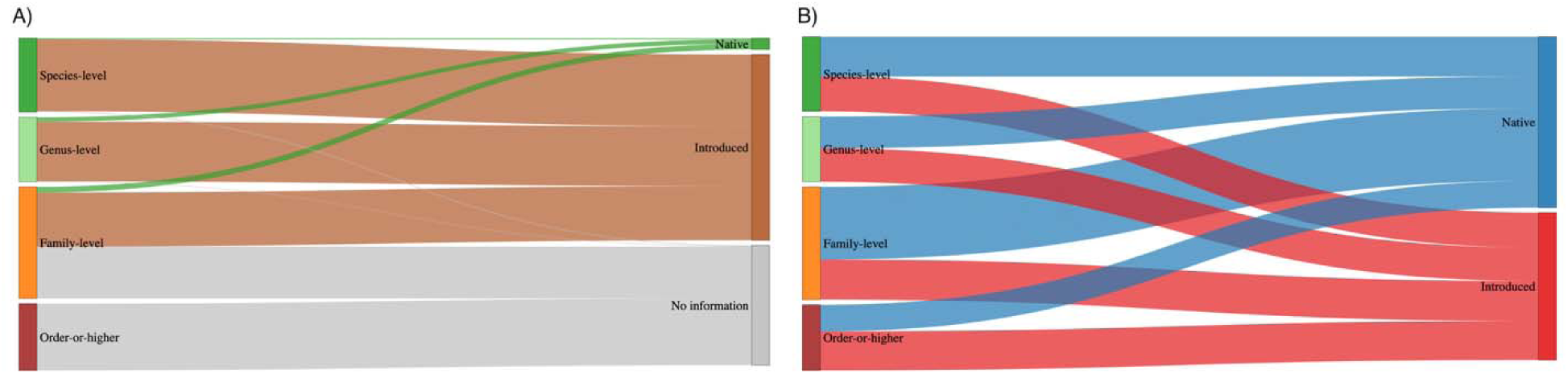
Sankey diagrams illustrating the increased utility of classifier predictions for ASV-level information over BLAST resolution by taxonomic level. A) Flow from BLAST-derived taxonomic resolution to invasion status inferred from reference-based assignments. Most status-resolved ASVs were assigned as introduced, whereas relatively few were linked to native taxa. B) Flow from taxonomic resolution to NIClassify status predictions, showing assignment of ASVs to native and introduced categories even when precise reference-based identification was unavailable. Together, the panels illustrate how classifier-based inference extends invasion-status assignment beyond the subset of ASVs with high-resolution BLAST matches.

**Table 1.** Comparison of invasion status assignments by classification method.

| Summary of Invasive Status Predictions |  |  |  |
| --- | --- | --- | --- |
| Prediction Method | Invasive Status |  |  |
|  | introduced | native | No information available |
| BOLD | 383 | 64 | 404 |
| IBIS | 433 | 22 | 396 |
| NCBI | 503 | 30 | 318 |
| NIClassify | 394 | 457 | 0 |
| Data obtained through Ibis geolocation API, BLAST mapping paired species lookup & NIClassify predictions |  |  |  |

### 3.2 Community differentiation across standardized O’hi’a gradients

NMDS ordinations based on Bray-Curtis dissimilarities showed broad overlap among low, mid, and high elevation samples, indicating that elevational separation in multivariate space was

modest at the island-wide scale (**Figure S3**). PERMANOVA detected limited variance explained by elevation alone (R² = 0.06, p = 0.40), whereas the combined model explained 37% of variation in community composition (R² = 0.37, p = 0.002). Multivariate dispersion differed among elevation bands (betadisper F = 5.18, p = 0.013). Sørensen analyses were congruent with Bray-Curtis results, with turnover averaging 0.73 ± 0.06 and varying across site and elevation (R² = 0.38, p = 0.001) under homogeneous dispersion (F = 0.78, p = 0.567). Bray-Curtis and Sørensen ordinations were strongly aligned (Procrustes RMSE = 0.10), and community ordinations were significantly associated with geographic distance from one each other (r = 0.79, p = 0.001). Pairwise Bray–Curtis dissimilarities also differed among grouping categories (Kruskal–Wallis χ² = 141.11, df = 6, p < 2 × 10 ¹; **Figure S4**). Median dissimilarity was lower within ridges (0.896) than between ridges (0.991), whereas differences by elevation band (within 0.986 vs between 0.990) were smaller. Geographic distance between samples was positively associated with Bray–Curtis community dissimilarity across samples (Mantel r = 0.284, p = 1 × 10, R² = 0.08; **Figure S5**).

#### 3.2.2 Richness and invasion patterns

Across standardized O’hi’a gradients, arthropod richness increased significantly with elevation (slope = 0.0466 ASVs m ¹, R² = 0.28, p = 0.017; **Figure S6**), although ridge-specific trends differed. Richness increased at Ka ala, declined at Poamoho, peaked at mid elevation on Mānana, and remained comparatively flat at Aiea and Schofield–Waikāne. Rainfall showed no consistent relationship with richness or the proportion of native ASVs across ridges and no significant partial effect once elevation was accounted for (**Figure S7**). The proportion of introduced ASVs declined significantly with elevation (Wald χ² = 9.95, p = 0.007), with mean introduced fractions of 0.64 ± 0.03 SE at low elevations, 0.54 ± 0.04 at mid elevations, and 0.47 ± 0.05 at high elevations (**Figure S8**). The proportion of introduced ASV sharing patterns were similarly narrow, with more than half of all ASVs detected on a single ridge and fewer than 10% shared among three or more ridges (**Figure S9).**

#### 3.2.3 Native and introduced subsets are both differentiated

When native and introduced subsets were ordinated separately, both showed strong differentiation in NMDS space (**Figure 3**). PERMANOVA detected significant compositional structure in both subsets (native: R² = 0.2554, F = 1.2860, p = 0.0054; introduced: R² = 0.3043, F = 1.6403, p = 0.0002), whereas multivariate dispersion did not differ between native and introduced assemblages (PERMDISP F = 0.1605, p = 0.5981).

**Figure 3.**
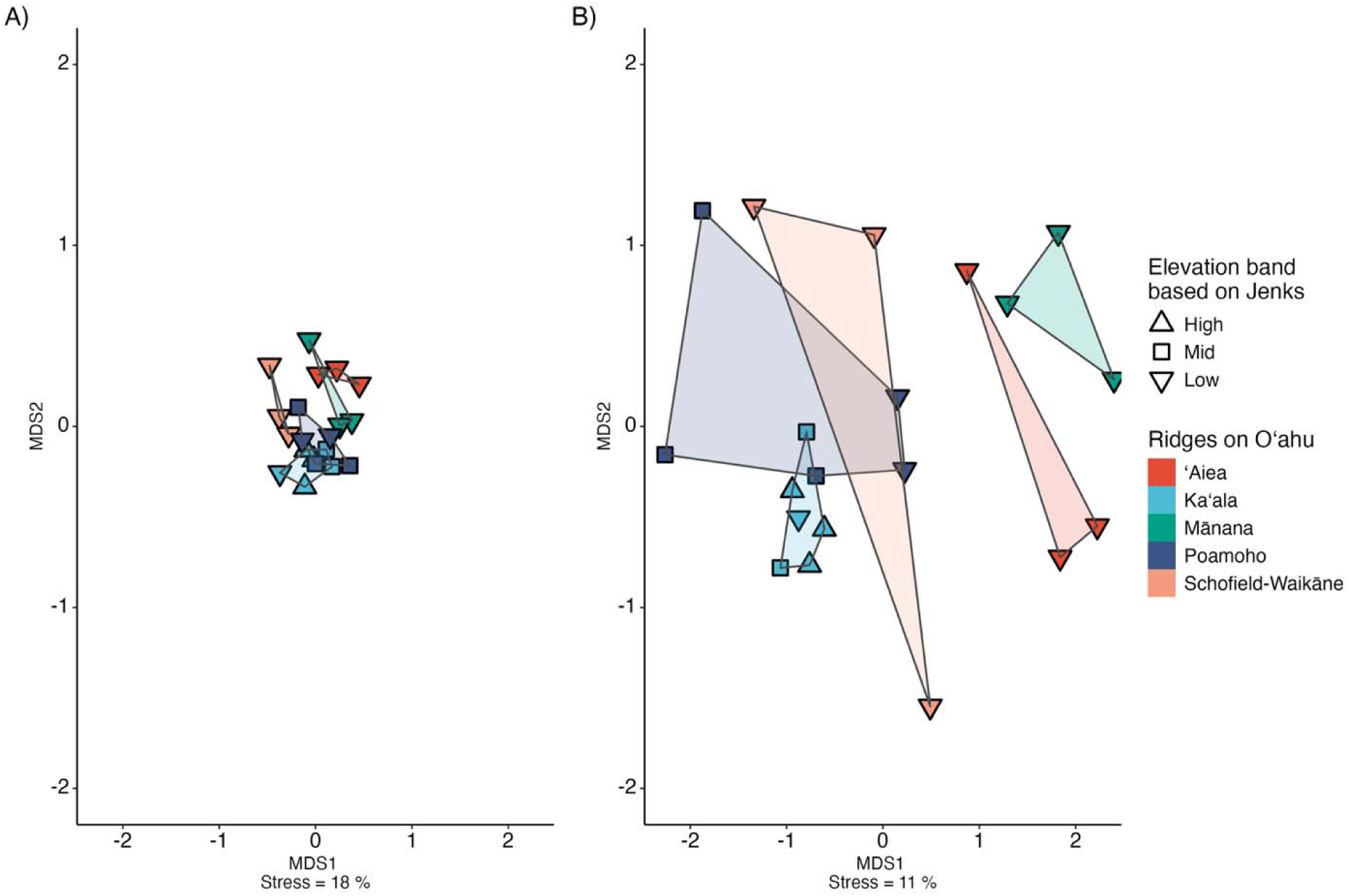
Native and introduced arthropod subsets remain compositionally differentiated across standardized O’hi’a gradients. NMDS ordinations based on Bray-Curtis dissimilarities for NMDS ordinations based on Bray-Curtis dissimilarities are shown separately for (A) native and (B) introduced ASV assemblages detected from O’hi’a across O’ahu ridges. Points represent biological samples, and polygons outline groupings in ordination space. PERMANOVA detected significant compositional structure in both subsets (native: R² = 0.255, F = 1.286, p = 0.0054; introduced: R² = 0.304, F = 1.640, p = 0.0002), whereas multivariate dispersion did not differ between subsets (PERMDISP: F = 0.1605, p = 0.5981).

### 3.3 Plant identity contributes to local species turnover

Native and invasive-status assignments for the arthropod ASV pool showed shared and unique combinations among O’hi’a, koa, and strawberry guava based on NIClassify (**Figure 4)**. Introduced ASV proportions varied among O’hi’a, koa, and strawberry guava at Aiea and Mānana (**Figure 5**). Based on locations, at Aiea, the mean percentage of introduced ASVs (± SE) was approximately 53% for O’hi’a, 43% for koa, and 48% for strawberry guava, and all pairwise contrasts were non-significant after Holm adjustment (p > 0.05). At Mānana, the corresponding means were approximately 66% for O’hi’a, 50% for koa, and 49% for strawberry guava. At Mānana, the proportion of introduced ASVs on O’hi’a exceeded that on strawberry guava (Holm-adjusted p < 0.05), whereas koa versus strawberry guava was non-significant. Across ridges, within-species comparisons were non-significant for O’hi’a, koa, and strawberry guava (all p > 0.05).

**Figure 4.**
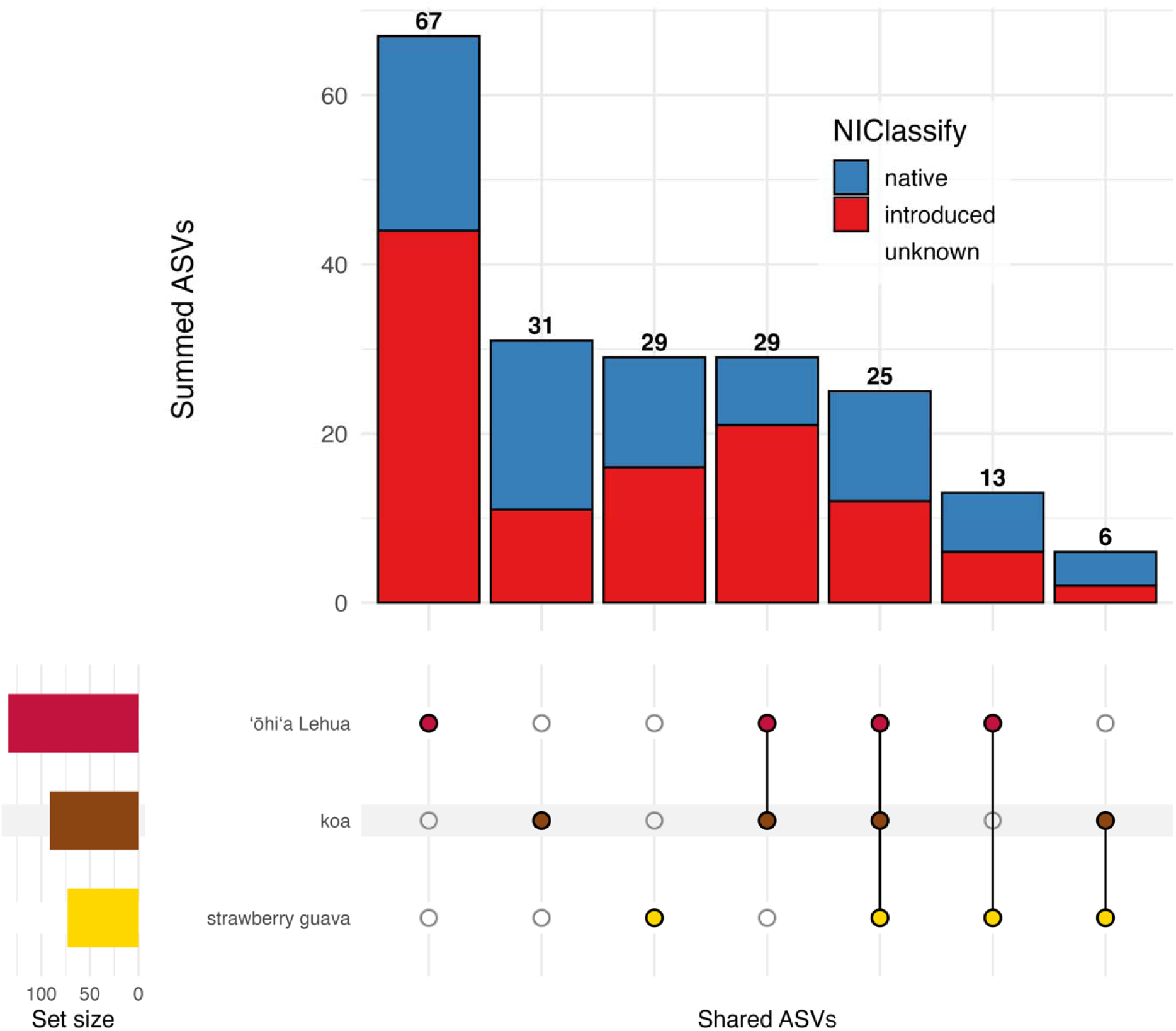
Overlap in arthropod ASVs among O’hi’a, koa, and strawberry guava based on NIClassify assignments.

**Figure 5.**
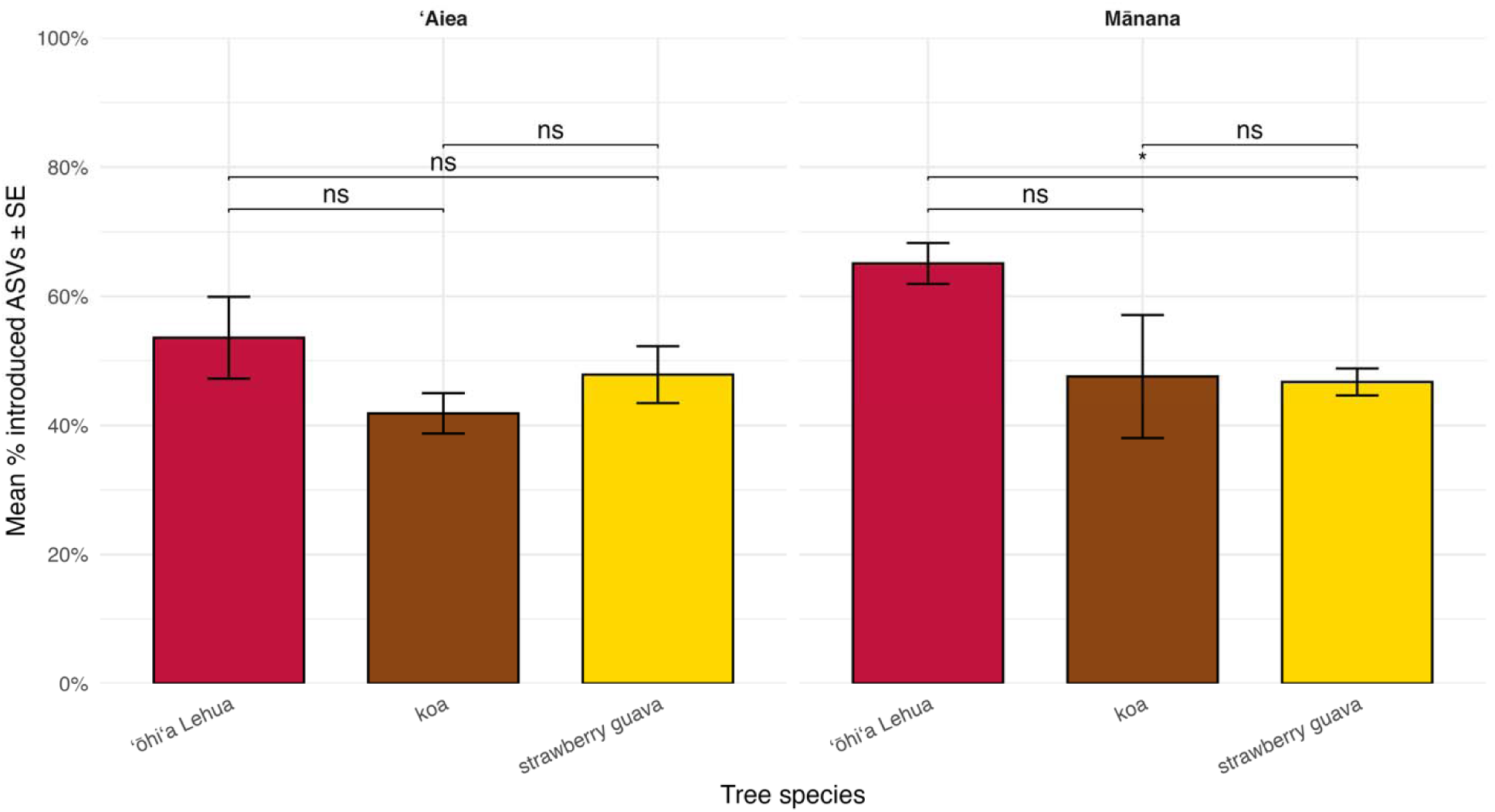
Ōhi a harbors more introduced taxa than koa and strawberry guava at Mānana Ridge. Bars show the mean proportion of introduced ASVs (± SE) for each tree species within each ridge. At Aiea, introduced fractions did not differ significantly among tree species. At Mānana, O’hi’a showed a higher proportion of introduced ASVs than strawberry guava, whereas the remaining pairwise contrasts were not significant after Holm correction. Colors indicate tree species: O’hi’a, koa, and strawberry guava.

### 3.4 A sharp community threshold occurs near 500 m elevation

Threshold Indicator Taxa Analysis (TITAN2) identified a clear elevation breakpoint in arthropod community composition at approximately 500 m. Presence–absence and abundance-weighted runs yielded overlapping change-point distributions, with bootstrap confidence intervals spanning 400 to 670 m. Using a purity and reliability cutoff of 0.95, 17 indicator ASVs were significant: five declining below approximately 450 m and twelve increasing between approximately 450 and 650 m. Increasing indicators represented multiple orders, including Amphipoda (n = 2), Collembola (n = 1), Isopoda (n = 1), Lepidoptera (n = 5), Coleoptera (n = 1), Hymenoptera (n = 1), and Trichoptera (n = 1) (**Figure 6**). Both native and introduced lineages were present among increasing indicators, whereas the declining group consisted mainly of low-elevation taxa. Relaxing the bootstrap reliability threshold to 0.90 increased the number of significant declining indicators and the proportion of introduced ASVs among them. The elevation of the community change point and the stability of its confidence range were consistent across analytical variants (**Figure S10**).

**Figure 6.**
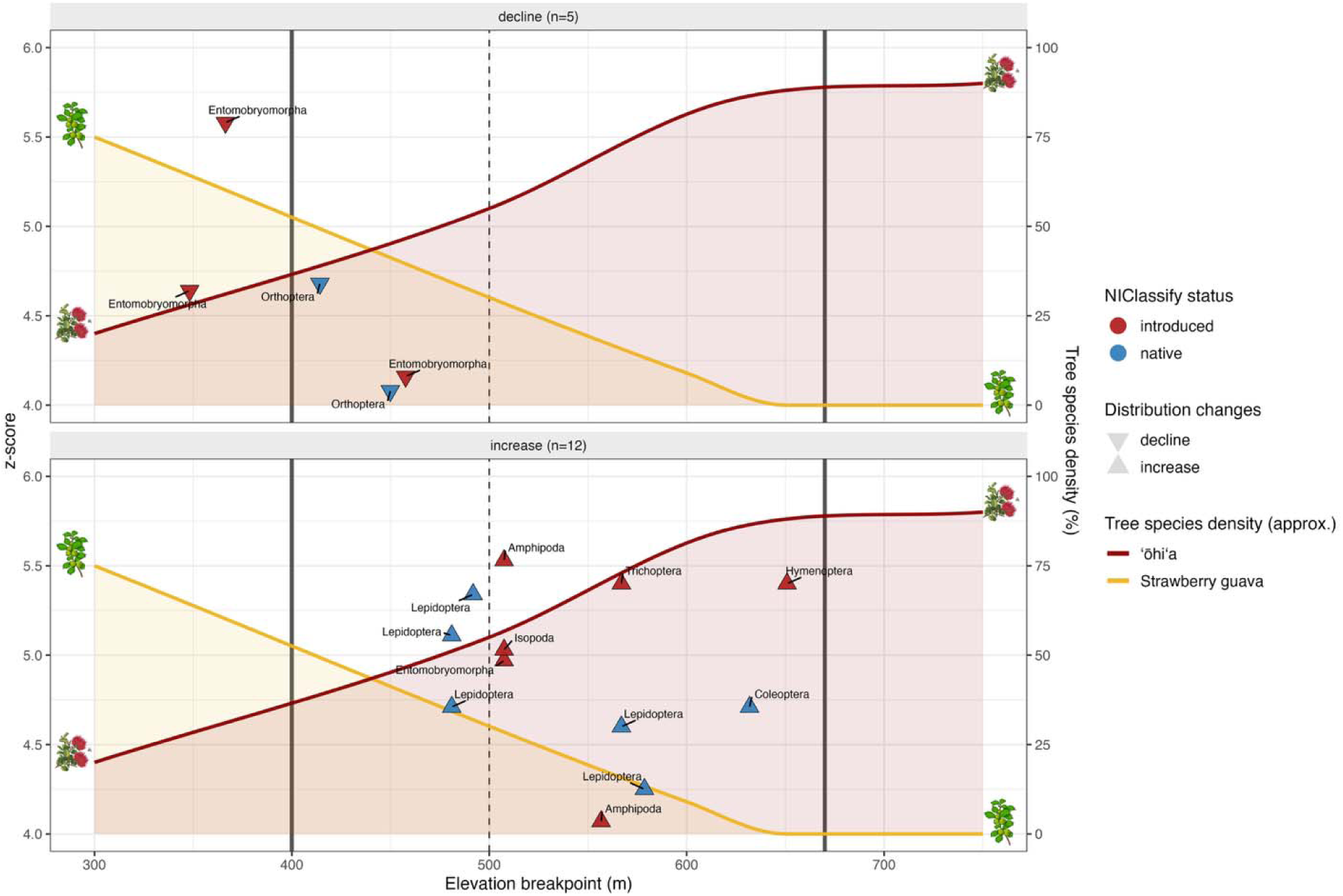
TITAN2 reveals a sharp turnover in arthropod communities at ∼500 m elevation. Per-ASV change points along O’ahu ridges identified by TITAN2 (purity ≥ 0.95, reliability ≥ 0.95). Triangles mark taxa that significantly decline (downward) or increase (upward) with elevation. Colors show NIClassify status (red = introduced, blue = native). The y-axis indicates indicator strength (z-score), while the secondary y-axis depicts approximate tree species density (% cover of O’hi’a and strawberry guava for sample sites on). The dashed vertical line marks the median community breakpoint (∼500 m), and the shaded area shows the 95 % bootstrap confidence interval (400–670 m). Declining indicators (n = 5) occur mostly below ∼450 m and are dominated by introduced springtails (Entomobryomorpha) with some native Orthoptera. Increasing indicators (n = 12) occur between ∼480–650 m and include both native groups (e.g., Lepidoptera, Coleoptera) and introduced taxa (e.g., Amphipoda, Isopoda, Hymenoptera).

## 4. Discussion

Arthropod communities detected from leaf-derived eDNA exhibited strong spatial differentiation across the O’ahu landscape, with substantial turnover among sampling regions and limited sharing of ASVs across sites. Such spatial structure is consistent with the fragmented topography of Hawaiian mountain systems, where ridges and valleys function as semi-isolated habitats that can promote lineage diversification and restricted dispersal (Lim et al., 2022). This dominant spatial signal constrains the ability to attribute community differences to single environmental gradients when sampling spans multiple regions. Accordingly, the present study emphasized analyses standardized on a single plant species across elevational gradients and local comparisons among co-occurring plant species, allowing tests of two specific hypotheses: (1) that if stronger native arthropod-native plant associations exist in less invaded, upper-elevation forests compared to more homogenization at lower elevations, the relative proportion of invasive arthropods should decrease with elevation, and (2) that plant identity, and especially plant origin (native or introduced), contributes additionally to local species turnover on shared slopes.

### 4.1 Elevation structures invasion signal in standardized O’hi’a communities

Elevation is a major organizing axis for native taxa in island ecosystems because it integrates climatic and habitat gradients that shape species distributions and community assembly (Körner, 2007; Steinbauer et al., 2016; Sundqvist et al., 2013) In Hawaiian forests, these transitions can occur over short geographic distances because climatic zonation is steep and compressed (Lim et al., 2022). Analyses standardized by sampling *M. polymorpha* across elevational gradients revealed shifts in arthropod community composition and invasion signal with increasing elevation. Richness increased upslope, the proportion of introduced ASVs declined, and community composition changed across a distinct mid-elevation transition.

The decline in introduced representation at higher elevations is consistent with patterns reported for many invasive taxa, which are often concentrated in warmer, more disturbed, lowland habitats and decline toward cooler, less accessible environments (Denslow, 2003; Loope et al., 2013; Pauchard et al., 2009). High-elevation forests on oceanic islands can therefore function as partial refugia for native biota, where climatic constraints, reduced propagule pressure, and lower human disturbance may limit the establishment of non-native species (Daehler, 2005; Vitousek et al., 2009). At the same time, the presence of both native and introduced indicator taxa above the transition zone from predominantly introduced to predominantly native species shows that this pattern reflects compositional turnover rather than simple exclusion of introduced lineages.

The observed increase in richness with elevation contrasts with the commonly reported decline in diversity toward higher elevations, although alternative patterns, including mid-elevation peaks or increases in less disturbed habitats, have been documented across taxa (Dolson & Kharouba, 2024; McCain & Grytnes, 2010; Rahbek, 1995). In our system, increasing richness with elevation coincided with a decline in introduced taxa and strong compositional turnover, suggesting that this pattern reflects a shift toward less invasion-dominated communities rather than a simple accumulation of species. However, the mechanisms underlying this pattern were not explicitly tested here.

The transition threshold from introduced to native arthropods (**Figure 6**) detected at approximately 500 m indicates that elevational change was not entirely gradual. Similar mid-elevation transitions have been reported in Hawaiian ecosystems and are often associated with shifts in rainfall regime, cloud influence, or vegetation structure (Giambelluca et al., 2013). The occurrence of both increasing and decreasing indicator taxa across this boundary is consistent with replacement among assemblages adapted to different environmental conditions rather than a unidirectional decline of low-elevation taxa. Such threshold responses are common in mountain systems where environmental change accumulates rapidly over short distances (Körner, 2007; Sundqvist et al., 2013).

### 4.2 Plant identity contributes local, context-dependent turnover

Our expectation was that plant identity shapes the compassions of arthropod communities, and that non-native plants would carry a higher proportion of invasive arthropods and have a homogenization effect on the geographic structure of arthropods overall. Indeed, comparisons among co-occurring plant species showed that plant identity contributed additional variation in arthropod communities, but effects on invasion signal were not consistent across ridges. Introduced ASV proportions differed among O’hi’a, koa, and strawberry guava at individual locations, yet no uniform pattern emerged in which non-native plants consistently supported more introduced taxa than the native species.

This absence of a simple native-versus-introduced dichotomy is consistent with the broader role of plant identity in shaping arthropod assemblages through differences in morphology, chemistry, phenology, and structural complexity (Novotny et al., 2002; Poelman et al., 2012; Southwood, 1961). Non-native plants can support both native and invasive arthropods, particularly when they share functional traits with resident vegetation or are integrated into local food webs (Heleno et al., 2009; Schirmel et al., 2016), likewise, native plants can serve as a resource of invasive arthropods. In Hawaiian forests, where many arthropod lineages use multiple plant species, community composition may therefore reflect a combination of plant traits and local species pools rather than plant origin alone (Kraft et al., 2015; Vellend, 2010).

The contrast between Aiea and Mānana suggests that plant identity effects operate within a broader environmental framework. Elevation, disturbance history, stand structure, surrounding vegetation, and propagule pressure can all influence colonization dynamics and community assembly, potentially overriding intrinsic differences among plant species. This interpretation is consistent with invasion ecology theory, which emphasizes habitat suitability, dispersal, and biotic interactions as key determinants of invasion outcomes rather than plant origin alone (Catford et al., 2009; Lockwood et al., 2005).

Because these comparisons were conducted on shared slopes where plant species co-occur, large-scale environmental differences were reduced and plant identity could be evaluated as a local factor. Within this framework, the results support an effect of plant identity on fine-scale community turnover, but not a general rule that native plants support predominantly native arthropod assemblages whereas invasive plants support introduced ones. Instead, invasion signal appears to emerge from the interaction between plant characteristics and local environmental context.

### 4.3 Invasion-status inference without complete taxonomic resolution

This study demonstrates the value of extracting invasion signal from DNA sequence-derived features rather than from taxonomic labels alone. Traditional approaches rely on assigning DNA sequences to named taxa and then consulting distribution databases, an approach that performs poorly in regions such as Hawai i, where endemic radiations are extensive but taxonomic coverage and sequence representation remain incomplete (Porter & Hajibabaei, 2018b; Weigand et al., 2019). In contrast, NIClassify produced status assignments for all ASVs without unresolved categories and recovered both native and introduced components within the community, including among sequences lacking precise matches in public databases. The contrast with BLAST-based ecological inference is particularly informative. Methods relying on nearest-reference matches tended to classify status-resolved ASVs overwhelmingly as introduced, reflecting well-known biases of genetic databases toward widespread, economically important, or medically relevant taxa rather than endemic diversity (Porter & Hajibabaei, 2018b, 2018a; Troudet et al., 2017; Weigand et al., 2019). By not requiring species-level naming, NIClassify avoids this asymmetry and allows community-level analyses that include poorly characterized lineages rather than excluding them as unresolved. This is particularly relevant for foliage-derived eDNA, which frequently captures rare, cryptic, juvenile, or transient taxa that lack reliable morphological identification or reference sequences (Deiner et al., 2017; Taberlet et al., 2012).

Importantly, the classifier-based approach does not replace taxonomic work but complements it. Status predictions derived from sequence features provide a functional categorization that can guide ecological interpretation and monitoring while formal identifications remain pending. As DNA reference libraries expand through continued barcoding and taxonomic revision, these assignments can be revisited and refined, but the present results demonstrate that invasion-relevant community patterns can be detected without complete species-level resolution. This capability is particularly valuable for early detection of invasive arthropods, where rapid assessment of community composition may be more informative than precise identification at initial stages of establishment (Darling & Mahon, 2011; Lodge et al., 2016; Simberloff et al., 2013).

More broadly, the ability to infer ecological status directly from DNA metabarcoding data expands the applicability of terrestrial eDNA for biodiversity assessment in data-poor regions. Oceanic islands, tropical ecosystems, and poorly studied taxa often share the combination of high diversity and limited reference coverage, constraining traditional taxonomy-dependent approaches (Gillespie et al. 2008). Methods that operate independently of species names therefore, offer a pragmatic pathway for extracting ecological information from complex communities while minimizing biases introduced by uneven taxonomic knowledge, and may be particularly useful for large-scale monitoring programs targeting cryptic or hyper-diverse groups (Deiner et al., 2017; Taberlet et al., 2012).

### 4.4 Limitations & implications for biodiversity monitoring and invasion detection

There are some clear limitations to our study. The first is sample size, and the extent to which differences might potentially be explained by incomplete sampling. Approaches such as rarefaction or asymptotic species richness estimators can tell us whether sample sizes are sufficient to represent the total species diversity in a community (Gotelli & Colwell, 2001) Despite this caveat, plant-derived eDNA recovered consistent differences in arthropod community composition across elevation and among co-occurring plant species, indicating that this approach can resolve ecologically relevant turnover in forest systems. Because sampling is non-destructive and logistically simple, it is well suited for repeated surveys in remote or protected habitats where conventional arthropod sampling can be difficult or disruptive (Basset et al., 2012; Stork et al., 2015). The ability to detect native and introduced components of the community from the same sampling framework is particularly relevant for invasion monitoring. Early detection remains central to invasive species management, yet conventional surveys often miss low-density, cryptic, or transient taxa during early stages of establishment (Lodge et al., 2016; Simberloff et al., 2013). Here, classifier-based status inference extended ecological interpretation beyond the subset of sequences that could be assigned confidently to named species, allowing invasion-related patterns to be assessed despite incomplete reference coverage.

Despite the potential of the overall approach, plant-derived eDNA has clear limitations. Detection depends on DNA deposition and persistence, and signals may include transient visitors rather than resident taxa. As a result, detections should not be interpreted as direct measures of abundance or strict plant association. In addition, ecological interpretation remains constrained when taxonomic resolution is low, even if invasion status can still be inferred. Continued improvement of DNA reference databases and integration with specimen-based sampling will therefore remain important for refining these inferences. Taken together, these results support plant-derived eDNA as a practical complement to traditional arthropod surveys and as a useful tool for tracking biodiversity change and invasion dynamics across heterogeneous forest landscapes.

## 5. Conclusions

Plant-derived eDNA combined with classifier-based status inference provides a practical framework for assessing arthropod communities in systems with incomplete taxonomic coverage. Because sampling can be standardized across sites, this approach is well suited for comparing community composition and invasion signal across heterogeneous island landscapes. In our study we were able to show the extent to which invasive arthropods have infiltrated native communities, across elevation gradients and host plants. Future work should test how stable these patterns remain through time and whether repeated sampling can detect shifts associated with climate change, disturbance, or new introductions. Integrating plant-derived eDNA with specimen-based surveys and improved reference databases will further strengthen taxonomic resolution and ecological. More broadly, the methods developed here may be transferable to other forest systems beyond Hawai i. Many regions with high biodiversity also lack comprehensive taxonomic coverage, limiting the effectiveness of traditional species-based monitoring. Approaches that infer ecological patterns directly from sequence data offer a practical pathway for studying community dynamics in such contexts while taxonomic knowledge continues to accumulate. As molecular tools and reference libraries improve, plant-derived eDNA is likely to become an increasingly important component of biodiversity assessment and conservation research.

## Data availability

Data is available for review and will be published after acceptance. http://datadryad.org/share/LINK_NOT_FOR_PUBLICATION/fBUWwDXxxaJjJ09qczZQTe-W3R9wA9gL80hiHG2OIVA

## Notes

### Competing Interest Statement

The authors have declared no competing interest.

### Funding

- SERDP: RC21-1034
- DFG: WE 7675/1-1

