## Supplementary material 1 for "Topography structures of arthropod communities revealed by leaf-derived environmental DNA on O’ahu, Hawai’i"

**Supplementary Materials for “Topography structures of arthropod communities revealed by leaf-derived environmental DNA on Oʻahu, Hawaiʻi” by Weber et al., 2026.**

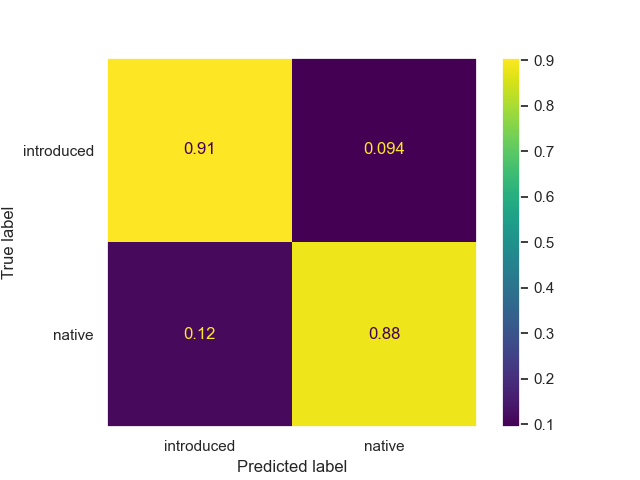

**Figure S1.** Cross-validation performance of the NIClassify invasion status classifier. Confusion matrix summarizing predictive performance of NIClassify during k-fold cross-validation using the labeled Hawaiian arthropod checklist dataset (n = 3,064 ASVs). Values represent the proportion of ASVs assigned to introduced or native categories relative to their known status. The classifier correctly identified 91% of introduced ASVs and 88% of native ASVs, indicating balanced predictive accuracy across invasion categories.

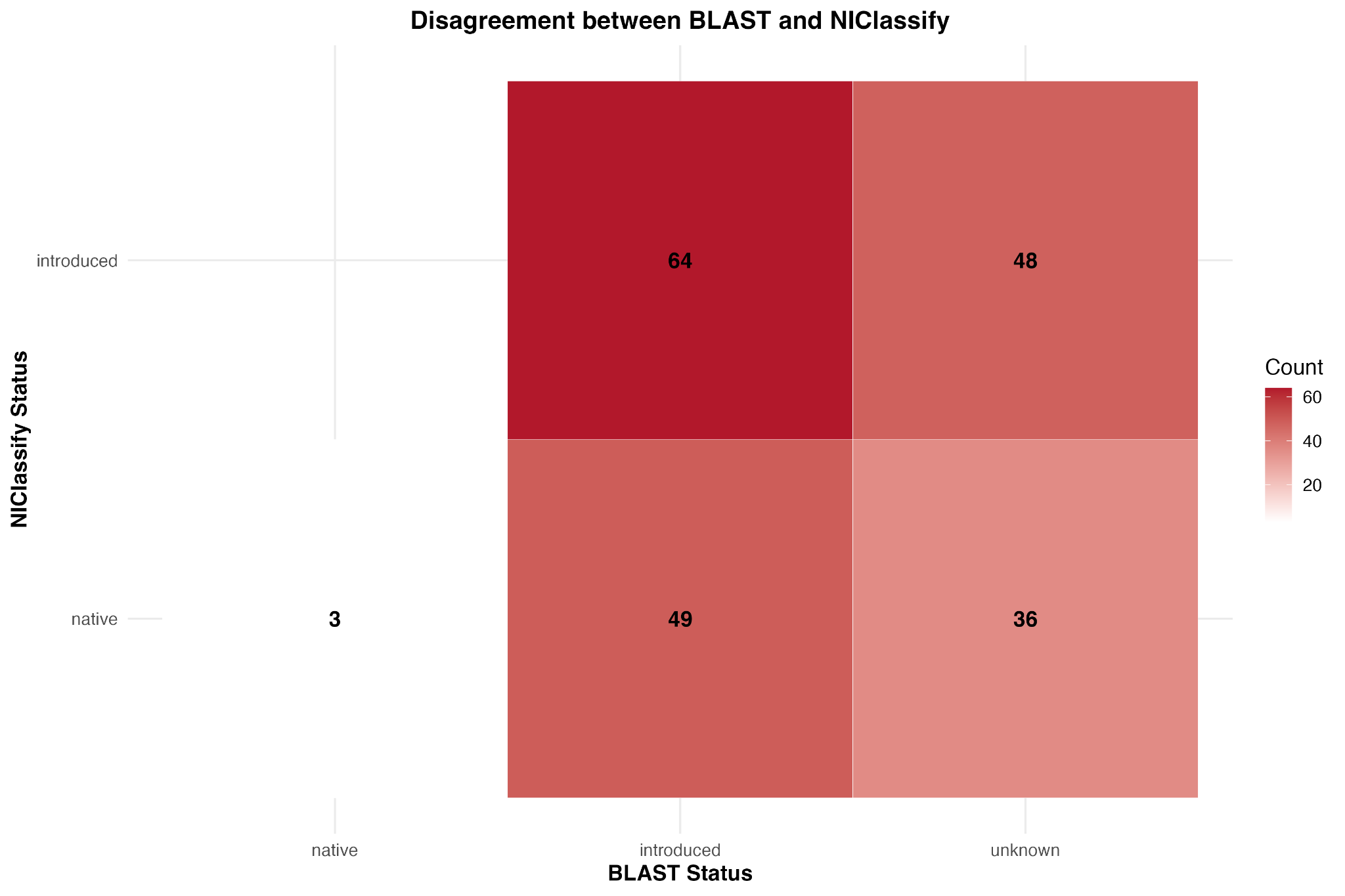

**Figure S2.** Disagreement in invasion status between BLAST-based inference and NIClassify predictions. Confusion matrix comparing invasion status inferred from BLAST-based ecological inference and NIClassify predictions for the 851 ASVs detected across Aiea and Manana ridges (ʻōhiʻa, koa, and strawberry guava). Counts represent the number of ASVs assigned to each invasion category under both methods. While both approaches identified a subset of ASVs as introduced, BLAST-based inference classified substantially more ASVs as introduced than NIClassify. Most disagreement occurred among ASVs labeled introduced by BLAST but predicted as native by NIClassify, illustrating systematic differences in invasion status inference under incomplete reference coverage.

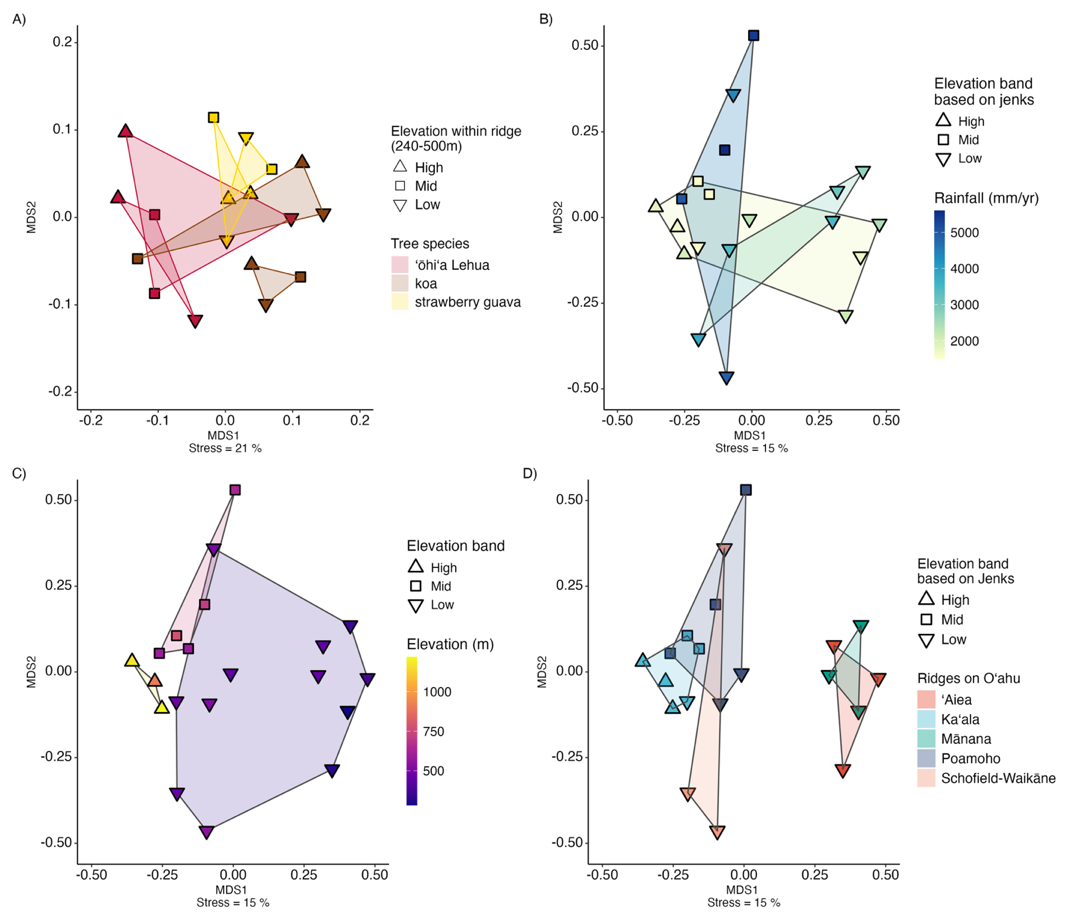

**Figure S3.** Ridge identity structures arthropod communities more strongly than elevation or rainfall. NMDS ordinations of arthropod communities from eDNA samples, based on presence–absence data. Symbols indicate elevation levels (triangles = high, squares = mid, inverted triangles = low). A) On ʻAiea and Mānana ridges, communities separate by tree (ʻōhiʻa, koa, strawberry guava), though stress is relatively high (21%). B–D Analyses restricted to ʻōhiʻa samples from all five ridges. (B) Grouping by annual rainfall bands shows no clear clustering. C) Assemblages grouped by elevation bands overlap broadly, with limited separation. D) Ridge identity produces the strongest signal, with samples from the same ridge forming distinct clusters (stress = 15%).

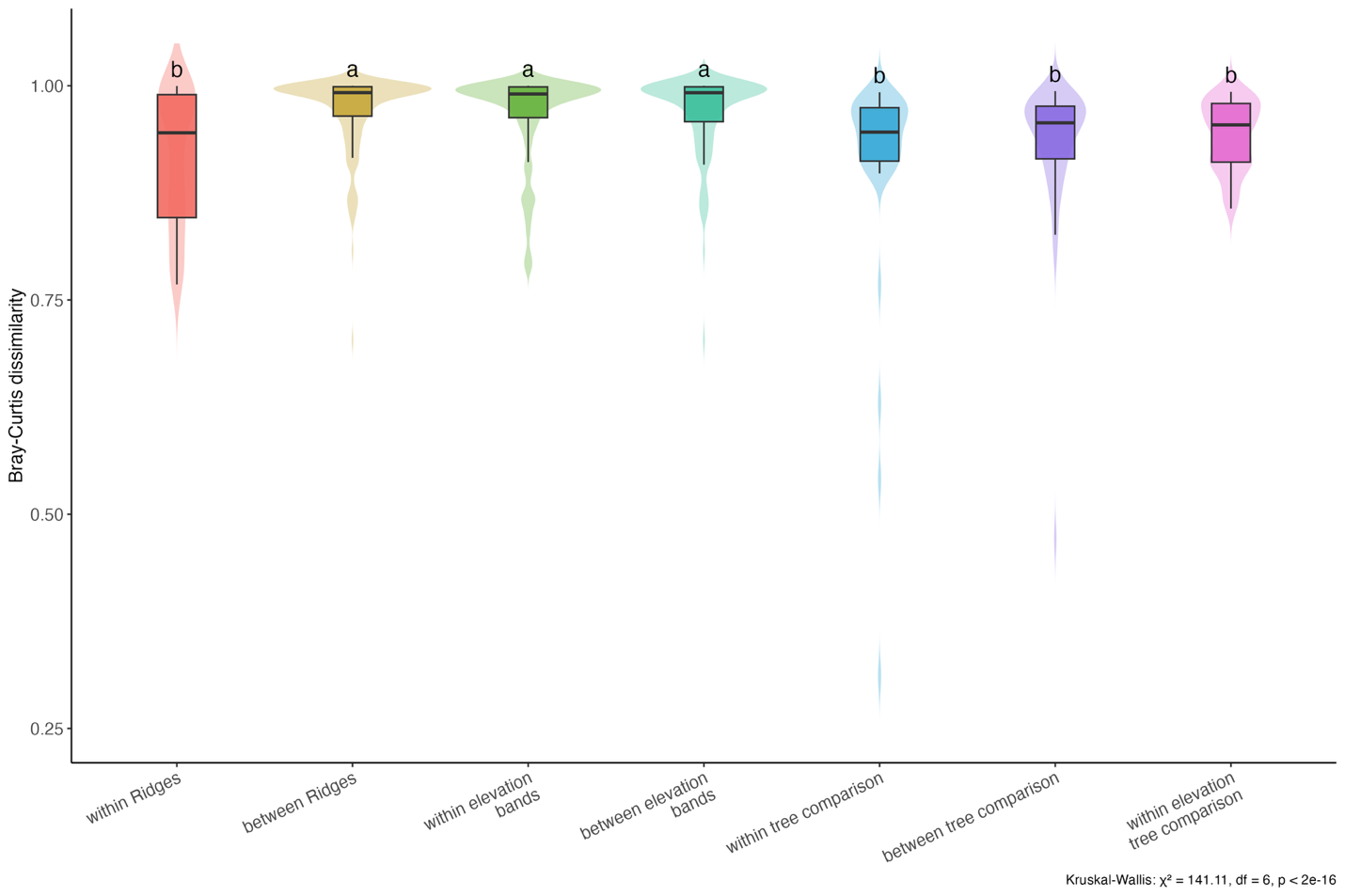

**Figure S4.** Community turnover across ridges, elevations, and plants. Distributions of Bray–Curtis dissimilarities between eDNA samples grouped by within- vs between-ridge, within- vs between-elevation, and within- vs between plant comparisons. Violin and boxplots show the distribution of pairwise dissimilarity values for each category. Community dissimilarities are substantially lower within ridges than between ridges (Kruskal–Wallis χ² = 141.11, df = 6, p < 2 × 10⁻¹⁶), indicating stronger spatial turnover among ridges than across elevation bands or species.

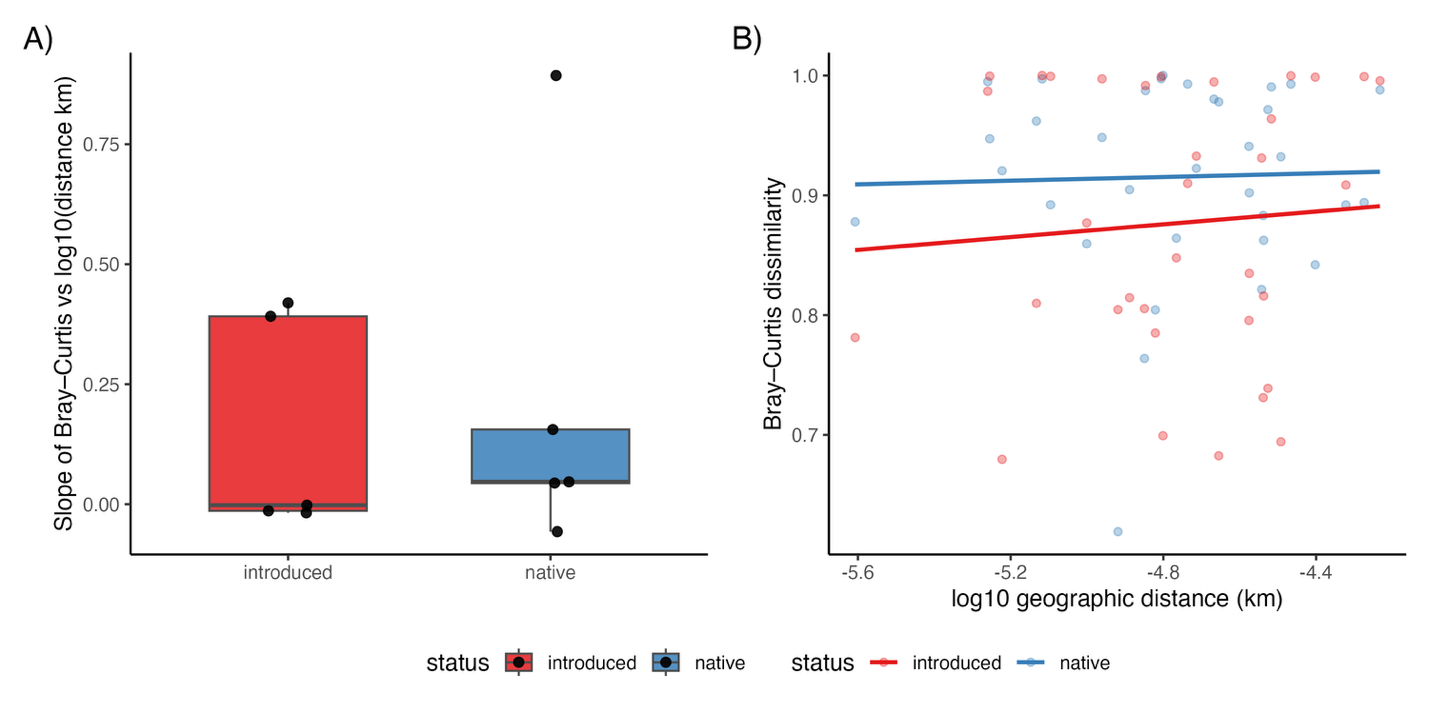

**Figure S5.** Community dissimilarity increases with geographic distance. The Mantel test indicates a weak but significant positive relationship between spatial distance and community dissimilarity, suggesting that geographic separation contributes modestly to arthropod community differentiation across the study system.

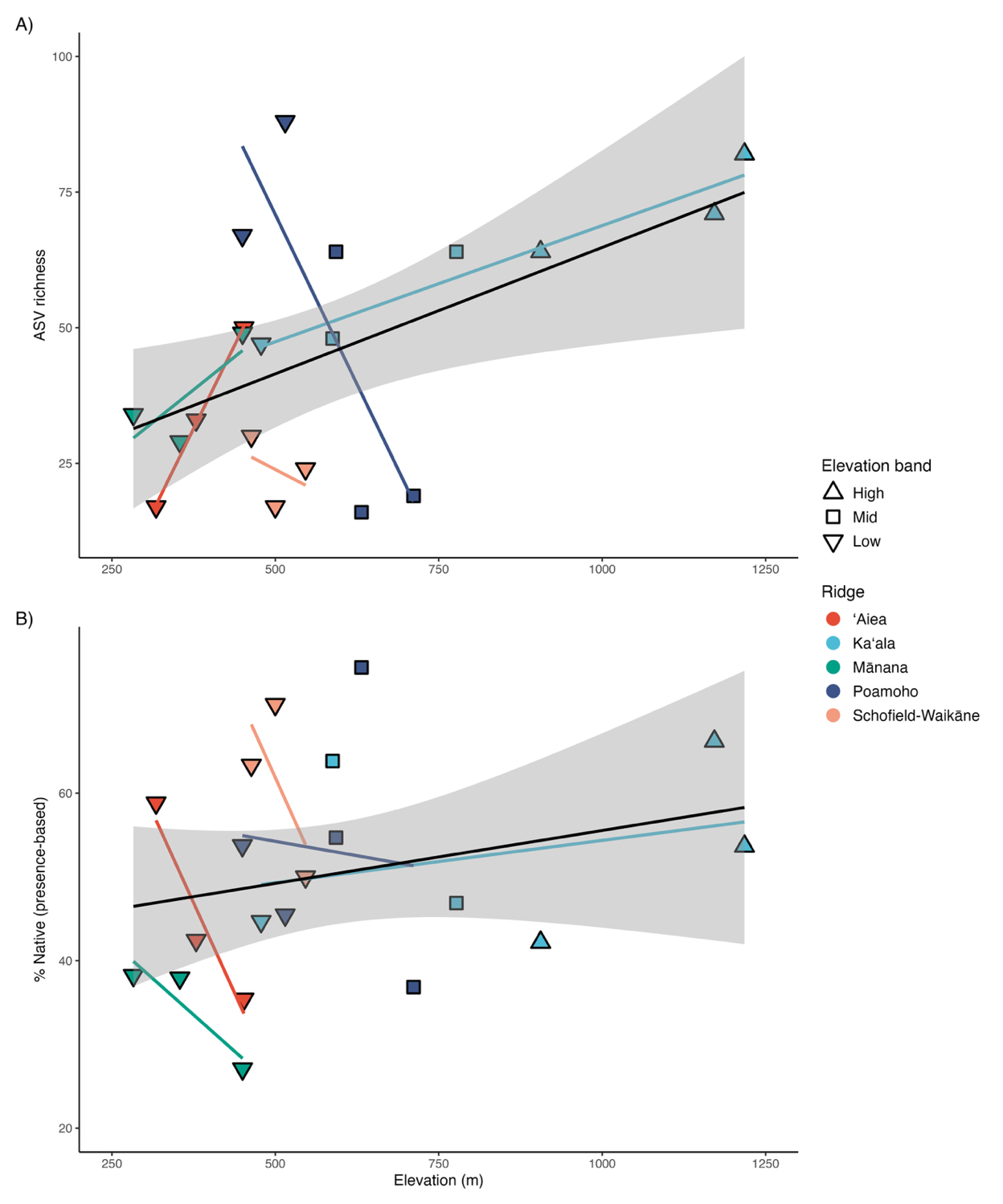

**Figure S6.** Arthropod richness increases weakly with elevation but varies strongly among ridges. Relationship between species richness (ASV counts) and elevation across all ridges. The pooled smoother (black line ± 95% CI) indicates a weak but positive trend, with higher elevation sites supporting on average more taxa than low-elevation sites. Colored lines show ridge-specific responses: richness rises with elevation at Kaʻala, declines at Poamoho, is hump-shaped at Mānana, and shows no clear pattern at ʻAiea or Schofield–Waikāne. These contrasting slopes highlight the dominance of site-level effects.

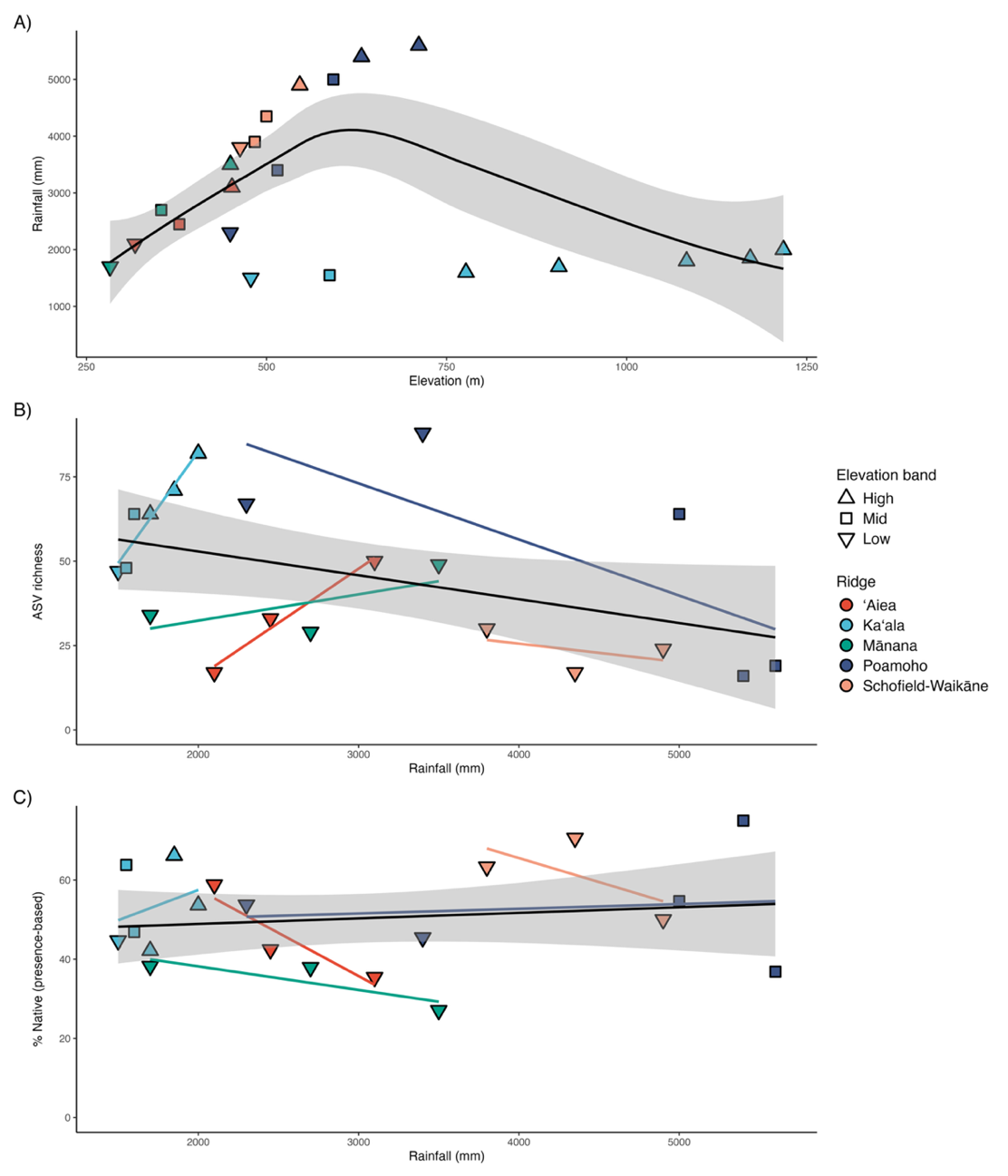
 **Figure S7.** Ridge-specific responses confirm weak explanatory power of rainfall. A) Relationship between elevation and mean annual precipitation across ridges. B) ASV richness plotted against precipitation. (C) Proportion of native ASVs plotted against precipitation. Black lines indicate overall trends across ridges, with colored lines showing ridge-specific responses. Together, the panels indicate that rainfall covaries with elevation but explains less variation in richness and nativeness than elevation-based analyses.

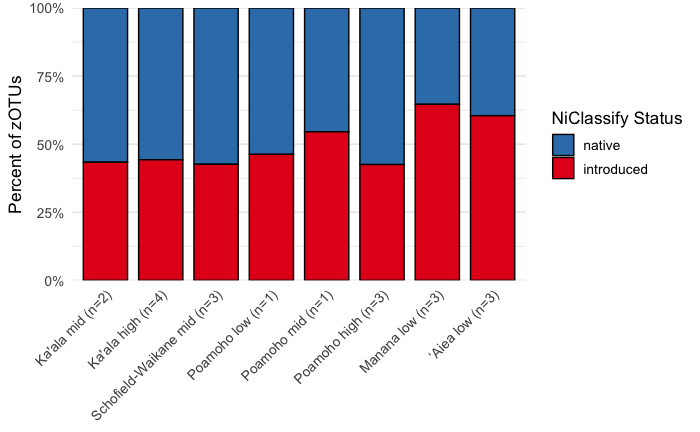
**Figure S8.** Elevation-associated invasion patterns in ʻōhiʻa-associated arthropod communities.
Stacked bars show the proportions of native and introduced ASVs across low, mid, and high elevation bands. Introduced fractions decline with increasing elevation, consistent with the binomial GEE analysis reported in the main text.

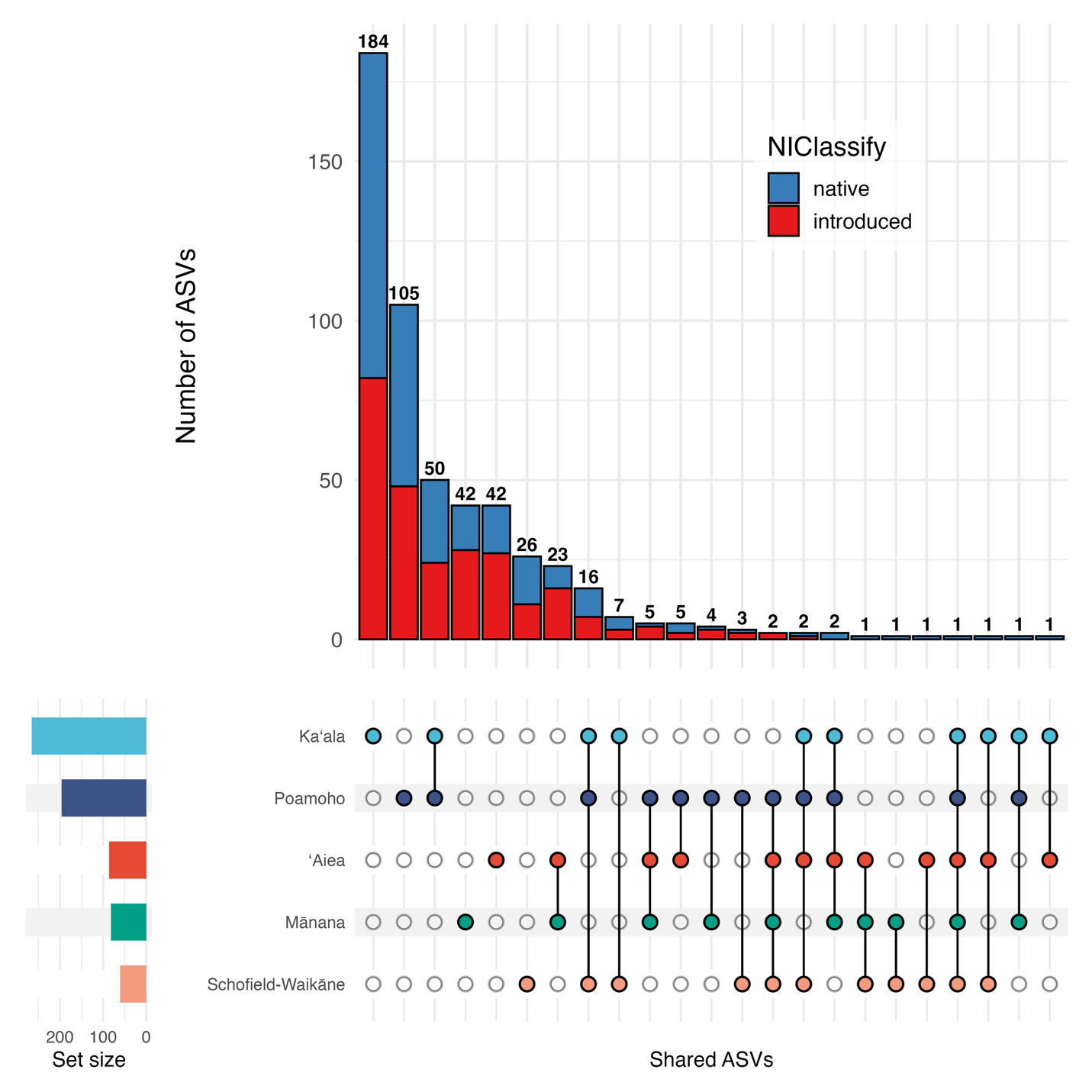

**Figure S9.** Most arthropod ASVs are ridge-restricted, consistent with an island-within-islands pattern. UpSet plot showing the distribution of ASVs across five Oʻahu ridges. On the left, horizontal bars give the total number of ASVs detected per ridge. In the central matrix, filled circles indicate which ridges contribute to each intersection set, and the vertical bars above show the number of ASVs shared by exactly those ridges. The tallest bars correspond to single-ridge intersections, demonstrating that most ASVs were unique to a single ridge, whereas only a small fraction were shared across multiple ridges. The stacked bar colors indicate the proportion of native (blue) versus introduced (red) ASVs within each intersection category, allowing direct comparison of native and non-native representation among unique and shared assemblages.

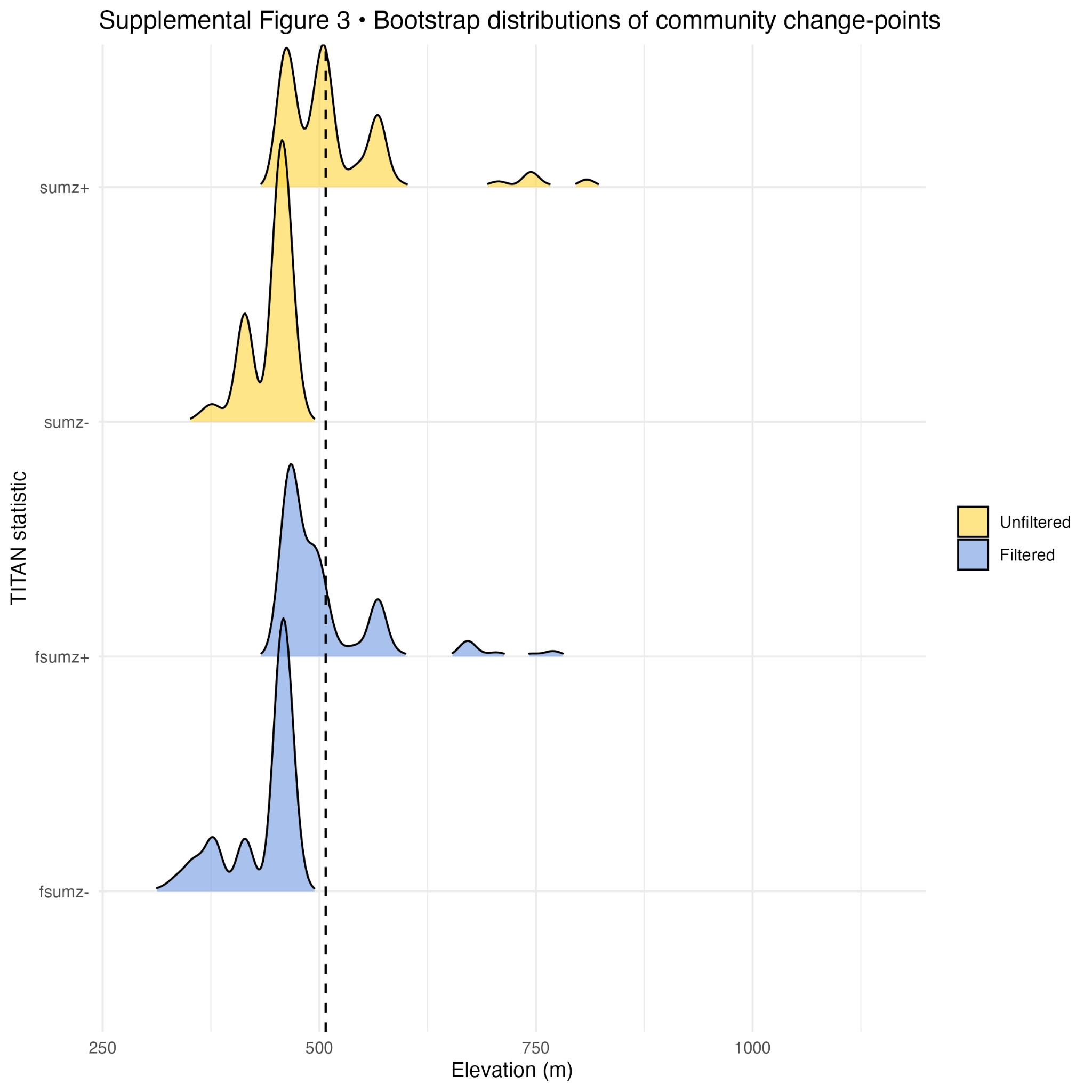

**Figure S10.** Bootstrap distributions of community change points along the elevation gradient. Density plots of bootstrap-estimated community breakpoints derived from TITAN analyses. Peaks indicate the most frequently inferred elevation thresholds where community composition shifts. Both unfiltered (red) and filtered (blue) analyses show a consistent concentration of change points near ~500 m elevation.

**Table S1**. Sampling locations, plant species, and the number of leaf-derived eDNA samples used for each location.

| Island | Ridge | Location | plant | Ridge elevation | Elevation (m) | Overall elevation | Trees sampled | Total sample |
| --- | --- | --- | --- | --- | --- | --- | --- | --- |
| Oahu | Ka ala | KA1 | *M. polymorpha* | Low | 478.2 | Low | 3 | 18 |
|  |  | KA2 |  | Mid | 587.7 | Mid | 3 |  |
|  |  | KA3 |  | High | 776.9 | Mid | 3 |  |
|  |  | KA4 |  | High | 905.9 | High | 3 |  |
|  |  | KA6 |  | High | 1171.7 | High | 3 |  |
|  |  | KA7 |  | High | 1217.7 | High | 3 |  |
|  | Manana | MA1 | *A. koa* | Low | 282.9 | Low | 3 | 9 |
|  |  | MA1 | *P. cattleianum* | Low | 282.9 | Low | 3 |  |
|  |  | MA1 | *M. polymorpha* | Low | 282.9 | Low | 3 |  |
|  | Manana | MA2 | *A. koa* | Mid | 353.9 | Low | 3 | 9 |
|  |  | MA2 | *P. cattleianum* | Mid | 353.9 | Low | 3 |  |
|  |  | MA2 | *M. polymorpha* | Mid | 353.9 | Low | 3 |  |
|  | Manana | MA3 | *A. koa* | High | 449.9 | Low | 3 | 9 |
|  |  | MA3 | *P. cattleianum* | High | 449.9 | Low | 3 |  |
|  |  | MA3 | *M. polymorpha* | High | 449.9 | Low | 3 |  |
|  | Poamoho | POA1 | *M. polymorpha* | Low | 449.6 | Low | 3 | 15 |
|  |  | POA2 |  | Mid | 515.1 | Mid | 3 |  |
|  |  | POA3 |  | Mid | 592.8 | Mid | 3 |  |
|  |  | POA4 |  | High | 631.9 | Mid | 3 |  |
|  |  | POA5 |  | High | 711.4 | Mid | 3 |  |
|  | Schofield-Waikane | KT0 | *M. polymorpha* | Low | 463.3 | Low | 3 | 9 |
|  |  | KT3 |  | Mid | 499.9 | Low | 3 |  |
|  |  | KT4 |  | High | 546.2 | Mid | 3 |  |
|  | Aiea | AL1 | *A. koa* | Low | 317.6 | Low | 3 | 9 |
|  |  | AL1 | *P. cattleianum* | Low | 317.6 | Low | 3 |  |
|  |  | AL1 | *M. polymorpha* | Low | 317.6 | Low | 3 |  |
|  | Aiea | AL2 | *A. koa* | Mid | 378.9 | Low | 3 | 9 |
|  |  | AL2 | *P. cattleianum* | Mid | 378.9 | Low | 3 |  |
|  |  | AL2 | *M. polymorpha* | Mid | 378.9 | Low | 3 |  |
|  | Aiea | AL3 | *A. koa* | High | 452 | Low | 3 | 9 |
|  |  | AL3 | *P. cattleianum* | High | 452 | Low | 3 |  |
|  |  | AL3 | *M. polymorpha* | High | 452 | Low | 3 |  |
